# Computational fluid dynamic analysis reveals the underlying physical forces playing a role in 3D multiplex brain organoid cultures

**DOI:** 10.1101/369082

**Authors:** Livia Goto-Silva, Nadia M. E. Ayad, Iasmin L. Herzog, Nilton P. Silva, Bernard Lamien, Helcio R. B. Orlande, Annie da Costa Souza, Sidarta Ribeiro, Michele Martins, Gilberto B. Domont, Magno Junqueira, Fernanda Tovar-Moll, Stevens K. Rehen

**Author notes:** Co-first authors (Livia Goto-Silva, Nadia M. E. Ayad).

## Abstract

Organoid cultivation in suspension culture requires agitation at low shear stress to allow for nutrient diffusion, which preserves tissue structure. Multiplex systems for organoid cultivation have been proposed, but whether they meet similar shear stress parameters as the regularly used spinner flask and its correlation with the successful generation of brain organoids, has not been determined. Herein, we used computational fluid dynamics (CFD) analysis to compare two multiplex culture conditions: steering plates on an orbital shaker and the use of a previously described bioreactor. The bioreactor had low speed and high shear stress regions that may affect cell aggregate growth, depending on volume, whereas the CFD parameters of the steering plates were closest to the parameters of the spinning flask. Our protocol improves the initial steps of the standard brain organoid formation, and organoids produced therefrom displayed regionalized brain structures, including retinal pigmented cells. Overall, we conclude that suspension culture on orbital steering plates is a cost-effective practical alternative to previously described platforms for the cultivation of brain organoids for research and multiplex testing.

**Highlights:**

- Improvements to organoid preparation protocol
- Multiplex suspension culture protocol successfully generate brain organoids
- Computational fluid dynamics (CFD) reveals emerging properties of suspension cultures
- CFD of steering plates is equivalent to that of spinner flask cultures

## 1. Introduction

Three-dimensional (3D) cerebral organoids generated from human pluripotent stem cells (hPSCs) are complex structures that partly reproduce fetal brain development *in vitro*, making them powerful tools for the study of human development and disease (Lancaster and Knoblich, 2014a). The self-organization that occurs during hPSC differentiation in cerebral organoids allows for the appearance of complex structures, including those recapitulating regions of the cerebral cortex, ventral forebrain, midbrain, hindbrain, hippocampus, and retina (Eiraku et al., 2011; Monzel et al., 2017; Quadrato et al., 2017). Several research groups have used this model to study the development of diseases such as microcephaly, lissencephaly (Bershteyn et al., 2017; Lancaster et al., 2013), and Zika infection (Garcez et al., 2016), as well as for drug testing(Dakic et al., 2017).

Organoids are grown in 3D suspension culture, which enables efficient nutrient delivery to 3D organized tissue. Historically, cerebral organoids have been cultured in spinner flasks (Lancaster and Knoblich, 2014b). These flasks have the advantage of providing a low-shear environment (Wang et al., 2013), which is important because hPSCs have been shown to be sensitive to shear stress(Nampe et al., 2017; Wang et al., 2013). However, spinner flasks have the disadvantage of requiring a high volume of cell culture media for cultivation, increasing the costs of experiments thus being limiting to drug testing and other multiplex experiments including comparison of multiple patients and controls. Recently, Qian et al. (2016)(Qian et al., 2016) proposed the use of a 3D-printed scalable mini-bioreactor, the SpinΩ, which would be cost effective and provide a feasible, reproducible platform for chemical compound testing. However, cultivation in the SpinΩ still requires an initial investment and the availability of 3D-printing equipment and other materials, which might make it infeasible for most laboratories. The use of orbital shaker plates described originally (Lancaster and Knoblich, 2014b) is a multiplex alternative to the often cost-prohibitive use of spinner flasks. However, whether the SpinΩ and orbital shaker plates provide the particle floating and nutrient mixing in a low-shear environment required to support organoid growth has not been addressed.

Here, we applied computational fluid dynamics (CFD) simulations to study shear stress and fluid flow in orbital shaker plates and the SpinΩ. Additionally, we developed an improved protocol to support the initial steps of organoid development (static phase), including embryoid body (EB) formation and compared suspension cultures in the SpinΩ bioreactor and orbital shaker plates (Lancaster and Knoblich, 2014b) in the initial 30-day cultivation period

## 2. Experimental Procedures

### 2.1. Pluripotent stem cell culture

The human induced pluripotent stem cells (iPSCs) used in this work are described in Supp. Table 1. GM23279 cell line from the NIGMS Human Genetic Cell Repository were obtained and certified by the Coriell cell repository; the remaining cell lines were generated in house. The iPSCs were maintained in six-well plates coated with Geltrex in mTeSR^®^ medium (StemCell Technologies, Canada). Cells were either passaged manually or with 0.15 mM EDTA through passage 48.

### 2.2. Culture of brain organoids

The method used to produce cerebral organoids was based on a previously published protocol (Lancaster and Knoblich, 2014b). Briefly, iPSC colonies grown in six-well plates were dissociated with 1 ml Accutase Cell Detachment Solution (MPBio, USA) for 4 min at 37°C, and 1 ml phosphate-buffered saline (PBS; LGC Biotechnology, USA) was then added. The resulting solution was transferred to a 15-ml conical tube, and 20 μl 10 mM Rho-kinase inhibitor (ROCKi, Y27632; Merck Millipore, USA) was added before centrifugation (Horiguchi et al., 2014), to obtain a final concentration of 10 μM. Cells were counted in a hemocytometer and centrifuged at 300 × *g* for 4 min. Cells were plated in hESC medium containing 50 μM ROCKi and 4 ng/ml b-FGF. hESC medium contained 20% knockout serum replacement (Life Technologies), 3% ESC-quality fetal bovine serum (Thermo Fisher Scientific, USA), 1% GlutaMAX (Life Technologies, Canada), 1% minimum essential medium non-essential amino acids (MEM-NEAAs; Life Technologies), 0.7% 2-mercaptoethanol, and 1% penicillin-streptomycin (P/S; Life Technologies), as described in Lancaster and Knoblich (2014) (Lancaster and Knoblich, 2014b). We used 9,000 cells/well of a 96-well plate, which has been demonstrated to lead to efficient organoid formation under the cultivation conditions described (Lancaster et al., 2017). The 96-well plates were centrifuged for 1 min at 300 × *g* to improve initial EB aggregation (Hookway et al., 2016) (Supp. Fig. 1b). After day 1, embryoid bodies (EBs) were cultured as described by Lancaster and Knoblich (2014) (Lancaster and Knoblich, 2014b). The medium was changed every 48 h after plating for 6 days. On day 6, EBs were transferred to 24-well ultra-low-attachment culture plates (one/well) containing 0.5 ml neuroinduction medium [1% N_2_ supplement (Gibco), 1% GlutaMAX (Life Technologies), 1% MEM-NEAAs, 1% P/S, and 1 μg/ml heparin in DMEM/F12 (Life Technologies). After 4 days (day 10), organoids were coated with Matrigel similarly as described by Sartore et al. (2017) (Sartore et al., 2017) in a 60-mm non-adherent tissue culture plate; six organoids were placed in 3 ml diluted Matrigel and incubated for 1 h at 37°C under 5% CO_2_. The coated organoids were then returned to the 24-well ultra-low-attachment plates with 0.5 ml neurodifferentiation medium with no vitamin A (50% neurobasal medium, 0.5% N_2_, 1% B_27_ supplement without vitamin A, 1:100 2-mercapto ethanol, 0.5% MEM-NEAA, 1% GlutaMAX, and 1:100 P/S in DMEM/F12) and left for 4 days in static culture. Subsequently, cerebral organoids were grown in suspension using two different platforms: 1) steering plates on a standard orbital shaker (six-well culture plates), agitated at 90 rpm [as proposed by Lancaster and Knoblich (2014)(Lancaster and Knoblich, 2014b)]; and 2) SpinΩ system developed by Qian et al. (Qian et al., 2016), which was 3D printed by the company DelthaThinkers using the blueprints provided in the manuscript and coupled to 12-well culture plates, agitated at 60 rpm. In both cases, 10 organoids were placed in 3 ml neurodifferentiation medium with vitamin A (day 14). The medium was changed weekly until day 60 of culture. They were imaged with an EVOS cell imaging system (Thermo Fisher Scientific) in brightfield. The area, diameter, and circularity of individual cerebral organoids were quantified using a custom macro in ImageJ.

### 2.3. Computational fluid dynamics simulation

CFD simulations were performed for the flows imposed by the SpinΩ impeller and the orbital shaker using the finite element commercial code COMSOL Multiphysics^®^. The rotational speed used for the SpinΩ was 60 rpm and that used for the orbital shaker was 90 rpm, with consideration of a 9.5-mm radius. The geometry and finite element meshes used in the simulations involved 3 ml suspension culture media for both mixing techniques (Supp. Fig. 2). The finite element meshes used for the bioreactor and 6 well plates simulations contained 390,000 and 125,000 elements, respectively. The numbers of finite elements and time steps used in the simulations were selected after grid refinement analyses; the differences between the velocity components computed with the meshes used in this work and with less refined meshes were less than 3% at selected points in the fluid domains. Although the mesh used for the well on the 6 well plate was practically uniform because regions of large gradients may occur at different positions during the periodic movement imposed externally by the plate (Supp. Fig. 2d and 3), the regions of largest gradients occurred at the edge of the impeller and on the surface of the rotating shaft in the case of the bioreactor, where the mesh was then refined (Fig. 2b, Supp. Fig. 2b). The simulations were performed with the *κ-ω* turbulence model for the SpinΩ. A laminar two-phase model, validated with the experimental results of Salek et al. (2012)(Salek et al., 2012), was used for the steering plates on the orbital shaker.

CFD simulation was performed with water as the working fluid and the assumption of Newtonian behavior. The flows were considered to be isothermal, after experimental evidence demonstrated that the work imposed by the impeller in the bioreactor was negligible (temperature variations over 24 h were less than 0.2°C; data not shown). Analysis of the orbital shaker required that transient states were simulated until a quasi–steady-state regime was reached, when the flow became periodic. Liquid flow was analyzed 0.5, 14, and 15 s after the start of movement

### 2.4. Histology and immunofluorescence

Cerebral organoids were fixed in 4% paraformaldehyde, incubated sequentially in sucrose solutions (10%, 20%, and 30%) prepared in PBS, embedded in optimal cutting temperature compound, and frozen in liquid nitrogen. The organoids were sectioned (20-μm thickness) with a cryostat (Leica Biosystems, Germany). Immunofluorescence was performed using the following primary antibodies: rabbit anti-nestin (RA22125, 1:500; Neuromics, USA), rabbit anti-PAX6 (42-6600, 1:100; Thermofisher Scientific), rabbit anti-TBR2 (AB2283,1:200; Millipore), mouse anti-MAP2 (M1406, 1:300; Sigma-Aldrich, USA), rabbit anti-FOXG1 (ab18259, 1:1,000; Abcam, UK), rabbit anti–islet-1 (ab20670; 1:1,000; Abcam), rabbit anti–OTX-2 (ab21990, 1:200; Abcam), mouse anti– glycogen synthetase (610518, 1:500; BD), and rabbit anti-PH3 (06-570, 1:500; Millipore). The following secondary antibodies were used: Alexa Fluor 488 goat anti-mouse (A11001, 1:500; Invitrogen, Canada) and Alexa Fluor 546 goat anti-rabbit (A11010, 1:500; Invitrogen). 4′,6-Diamidino-2-phenylindole (1 mg/ml) was used for nucleus staining. Images were acquired using a Leica TCS SP8 confocal microscope.

### 2.5. Proteomic analysis

Two independent pools of four organoids each were used in the experiments. Protein digestion, peptide fractionation, mass spectrometric analysis, and raw data processing was performed as described by Murillo et al. (2017) **(Murillo et al., 2017)**. Gene enrichment analysis was performed using the DAVID Bioinformatics Database (https://david.ncifcrf.gov/summary.jsp).

### 2.6. Statistical analysis

Statistical testing was performed using two-tailed t-test with GraphPad Prism 6 software. Statistical significance was defined as p<0.05 unless otherwise stated in figure legends. Correlation analysis was done comparing the R square of a non-linear fit (Exponential fit in Fig. 1c and a Gaussian fit in Fig. 1e) for the two conditions, SpinΩ and orbital shaker.

**Figure 1.**
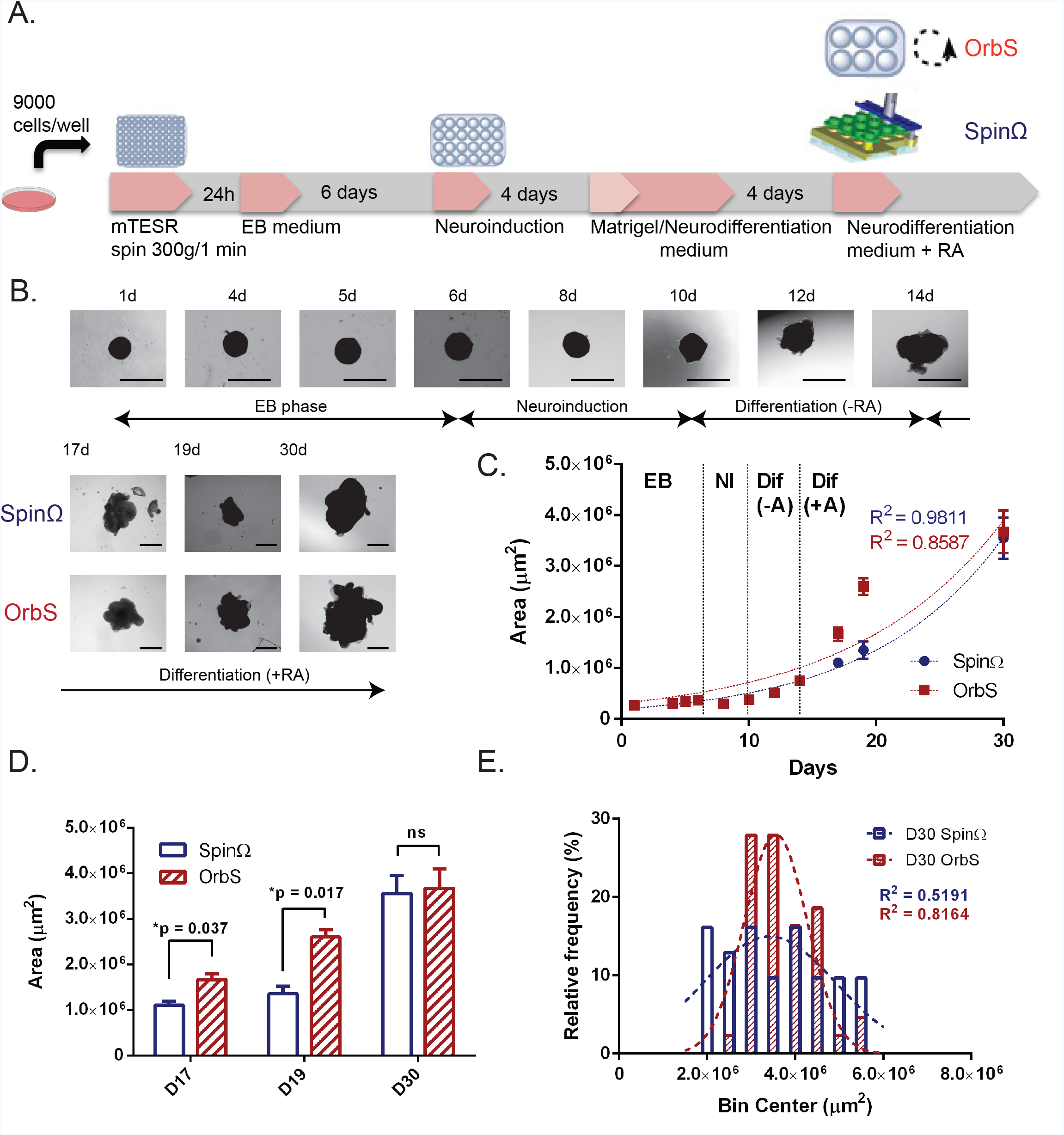
Growth curves, size distribution, and morphology obtained with the orbital shaker and SpinΩ bioreactor. **a**. Workflow for brain organoid preparation. **b**. Organoid morphology on selected days of the 30-day culture period, with arrows indicating the length of exposure to each medium condition. On day 14, organoids were divided into growth in the orbital shaker (red) and in the SpinΩ (blue). Scale bar = 1,000 μm **c**. Area growth curves (μm^2^) from day 1 to day 30, from n = 2 independent tests. Each test replicate contained at least 12 individual brain organoids. Lines represent exponential curve fit for the SpinΩ (blue) and orbital shaker (red), with correlation coefficients for each curve displayed on the graph. **d**. Comparison of area growth between the SpinΩ and shaker groups \**p* < 0.05. **e.** Histographic analysis of organoids grown in the SpinΩ and orbital shaker on day 30 (area in μm^2^) showing relative frequencies in terms of the percentage of automatically binned size. Data were pooled from two independent tests, *n* = 31 for the SpinΩ and *n* = 43 for the orbital shaker.

## 3. Results

### 3.1. Improved embryoid body (EB) formation and analysis of organoid growth

To improve the first step of EB formation, we introduced some variations to the protocol published by Lancaster and Knoblich (2014) (Lancaster and Knoblich, 2014b). The experimental flow is depicted in Fig. 1a. Changes in the protocol include the addition of Rho-associated protein kinase inhibitor (ROCKi) for cell survival at cell dissociation (Horiguchi et al., 2014) and post-plating centrifugation (Hookway et al., 2016). iPSCs cultivated in mTeSR1 medium derived from manual passages showed better EB formation compared with Ethylenediaminetetraacetic acid (EDTA)-passaged iPSCs (data not shown). Therefore, the cells were passaged manually before the EB formation step. Immediately after treatment for cell dissociation and before centrifugation, 10 μM ROCKi was added to the trituration solution. This step improved cell morphology after dissociation (Supp. Fig. 1a and b)(Horiguchi et al., 2014). The centrifugation step significantly improved the circularity of organoids on day 1 of growth (Supp. Fig. 1e), which was correlated with a significant increase in the observed areas of organoids in the two conditions (Supp. Fig. 1f). However, after 10 days, no significant difference was seen between specimens treated with and without centrifugation (Supp. Fig. 1f), suggesting that this potentially negative effect was temporary. During the EB stage, no significant growth was observed (Fig.1a, b) and the morphology of aggregates did not change (Fig. 1b). Growth during the neuroinduction stage also was not significant (Fig. 1a, b). In the neuroinduction stage, protrusions of developing organoids started to expand; these continued to grow over time (Fig. 1b) and formed neuroepithelium-like tissue (see also Fig. 3b). The pattern of organoid growth resembled an exponential curve (Fig. 1c), with R square values of 0.9811 for the SpinΩ and 0.8587 for the orbital shaker. Growth in the orbital shaker was initially more rapid than that in the SpinΩ (Fig. 1d), but no significant difference was observed at the 30-day timepoint. A histographic analysis of organoids grown in the SpinΩ and the orbital shaker at 30 days showed that the size distribution of organoids grown on the shaker more closely resembled a Gaussian fit (Fig. 1e), suggesting more homogeneity in the shaker.

### 3.2. The orbital shaker delivered higher velocity fields, but less shear stress, than the SpinΩ

The fluid dynamics in cultivation vessels have been shown to influence cell stemness, differentiation, and growth (for a review, see (Fridley et al., 2012)). Here, we analyzed the fluid dynamic conditions to which organoids were subjected to understand the differences in growth in the SpinΩ and the orbital shaker plates. Although some regions of the well were depleted of fluid at 0.5 s, just after the start of the plate movement, the fluid eventually covered the whole bottom surface of the well as the flow developed for the quasi–steady-state regime. The velocities were symmetrical at 14 and 15 s, due to the periodic flow created by the circular movement of the shaker (Supp. Fig. 3). Figure 2a presents the absolute velocity fields for the well on the orbital shaker at 15 s after the initiation of plate movement. The maximum absolute velocity reached with the 6 well plate on the orbital shaker during the quasi–steady-state regime was about 0.12 m/s. The shear stress field at 15 s is shown in Figure 2b. The maximum stress was about 0.045 Pa in the regions of maximum-velocity gradients near the walls (see also Fig. 2a). The magnitude of the shear stress was about 10^−2^ Pa in a large region of the fluid (Figure 2a).

**Figure 2.**
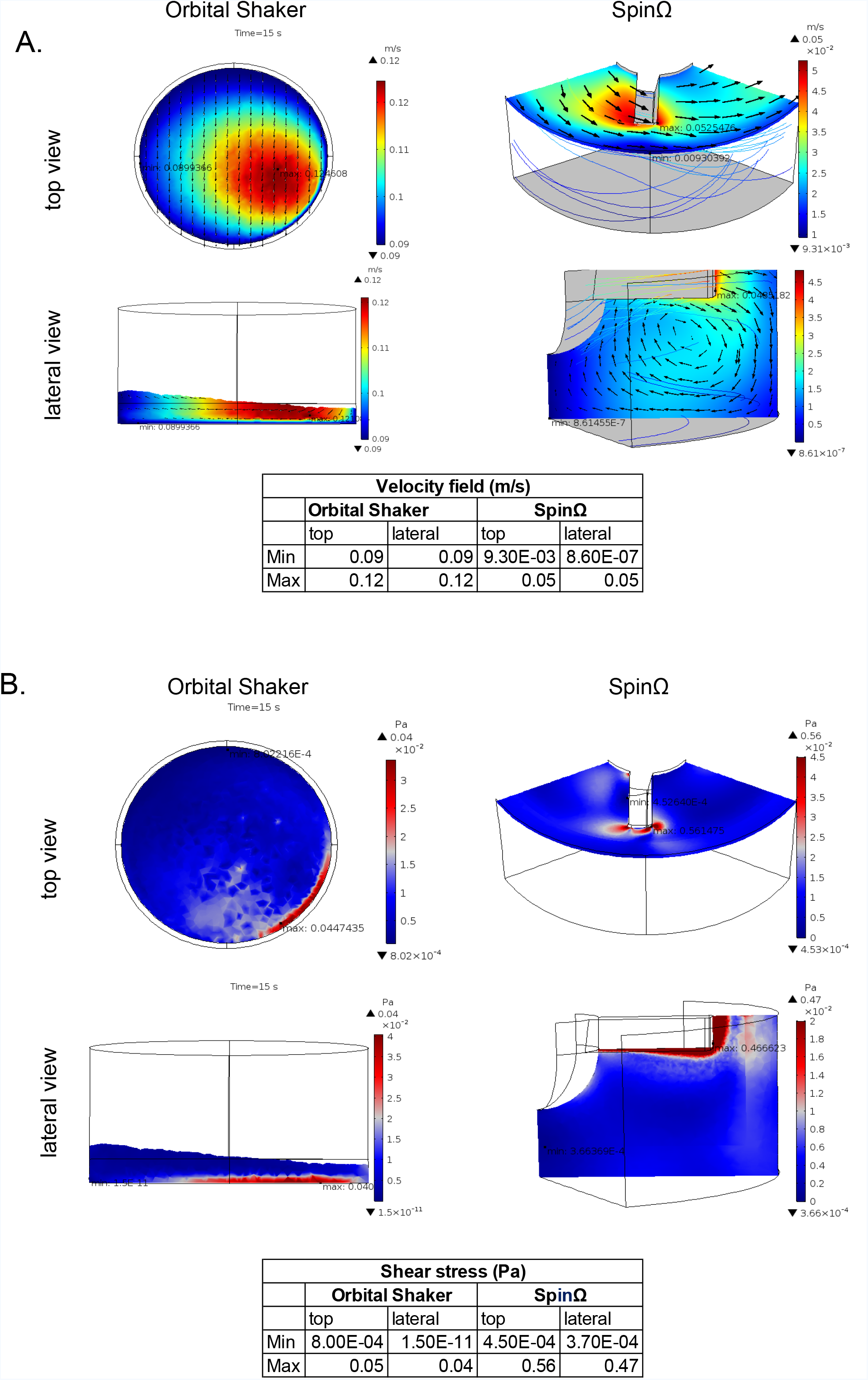
Fluid dynamic models in suspension culture. Computational fluid dynamic analysis of fluid velocity and shear stress for the SpinΩ and orbital shaker. **a.** Velocity fields and **b.** shear stress, top and lateral views for the SpinΩ and orbital shaker at 15 s after the start of movement. Minimum and maximum values are presented in the tables.

The SpinΩ analysis is presented for different plane cuts. The velocity and shear stress fields are presented in Figure 2b. The highest velocities occurred at the edge of the impeller, with values around 0.05 m/s (Fig. 2a). The velocities decreased with distance from the impeller and rotating shaft, being null at the well walls due to the non-slip conditions. In particular, lower velocities at the bottom of the well did not favor the mixture required for the enhanced growth of organoids. Our attempts to form aggregates from single cells in the SpinΩ created bodies with disparate sizes. Single cells accumulated in the low-speed area of the bioreactor and formed a large aggregate, while adjacent cells formed smaller bodies (Supp. Fig. 4). Large differences in the velocity fields implied differences in nutrient mixing, which could in part explain the delayed growth in the SpinΩ on days 17–19 (Fig. 1d) and the wide area distribution shown on the histogram at day 30 for SpinΩ (Fig. 1e).

The maximum shear stress was 0.56 Pa at the edge of the impeller due to the large difference between velocity gradients in this region (Fig. 2b). Shear stress in the bulk fluid was lesser, with magnitudes on the order of 10^−3^ to 10^−2^ Pa.

Velocity and shear stress are correlated parameters. In this study, however, the absolute velocity magnitudes were greater for the orbital shaker than for the SpinΩ, whereas shear stress values had the opposite pattern. These results can be explained by the large differences in velocity values at the region around the SpinΩ impeller and the rotating shaft increasing the shear of fluid in this area.

The gold standard for organoid protocols, a spinning bioreactor has been reported to sustain organoid growth in culture for more than 8 months (Lancaster and Knoblich, 2014b; Quadrato et al., 2017). Comparison with previous literature on the CFD of a spinning bioreactor (Wang et al., 2013) showed that shear stresses of the steering plates found on this study are of the same order of magnitude as those reported for the spinner bioreactor (Wang et al., 2013): maximum shear stress values for the spinner were 0.028 Pa at 40 rpm and 0.047 Pa at 75 rpm (Wang et al., 2013) (Supp. Table 2), and a maximum value of 0.045 Pa at 90 rpm was predicted for the orbital shaker (Supp. Table 2). The maximum shear stress of the SpinΩ (0.56 Pa) was one order of magnitude greater.

The velocity fields of the steering plates (maximum, 0.12 m/s) were of the same order of magnitude as those of the spinner bioreactor (maximum, 0.277 m/s), whereas those of the SpinΩ were lower (0.05 m/s) (Supp. Table 2).

Computational simulations indicated that the use of the 6 well plates on the orbital shaker was more suitable for the growth of organoids than was the use of the bioreactor. First, the SpinΩ has regions of low velocity at the bottom of the well, where fluid mixing is poor and particles deposition is likely to happen. Second, shear stress is lesser in the orbital shaker, which could be better for the preservation of organoid structures in long-term culture.

### 3.3. The SpinΩ reactor and orbital shaker derived structured organoids

We examined the maturation of organoids with a focus on the transition from predominantly neuroprogenitor stem cells to the development of neuroepithelial regions. Nestin staining of neural stem cells, performed at 10, 14, and 30 days of culture, showed similar decreases over time for the orbital shaker and SpinΩ treatments (Fig. 3a). These results are consistent with the start of differentiation of progenitor cells into neurons. At 30 days, the organoids had developed ventricular-like regions and neuroepithelium-like structures that were positive for MAP2 and TBR2 (Fig. 3b). MAP2 staining levels were similar in organoids cultivated in the orbital shaker and SpinΩ, but TBR2 staining levels were significantly stronger for those cultivated in the SpinΩ. As TBR2 labeled neuron progenitors in sub-ventricular zones, we examined whether cell proliferation was increased under our culture conditions through phospho-histone-3 staining on day 30. However, no difference in the number of proliferating cells was detected between the two conditions (Fig. 3c).

**Figure 3.**
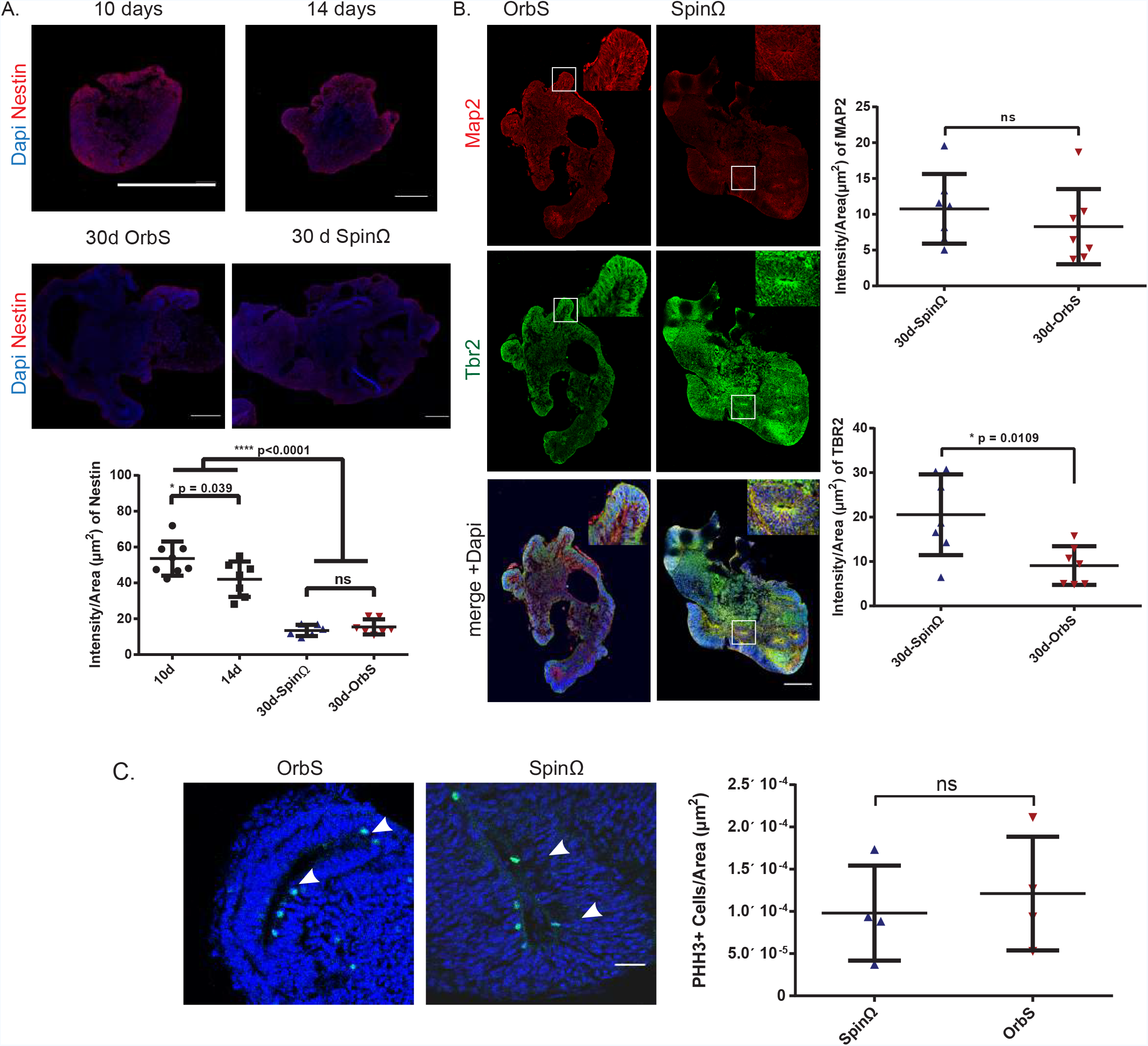
Neurogenesis in the early stages of organoid development. **a.** Top, representative nestin immunostaining of organoids after 10, 14, and 30 days in culture. Bars = 500 μm. Bottom, quantification of intensity of nestin staining normalized with area of organoid for 10, 14 and 30 days in culture for the different conditions. N= 8, 7, 7 and 6, respectively. **b.** Left, representative MAP2 and TBR2 staining of 30-day-old organoids. Bars = 500 μm. Right, quantification of intensity of MAP2 and TBR2 staining. N = 7 organoids for each condition. **c.** Left, representative phospho-histone H3 staining of ventricular-like regions of 30-day-old organoids. Bars = 100 μm. And right, corresponding quantification. N = 4 for each condition. For all staining, organoids were collected from 2 independent experiments. Mean + SD for all quantifications.

### 3.4. Organoids generated in suspension cultures presented markers for distinct brain regions

Organoids grown in the SpinΩ and orbital shaker displayed very similar morphology and developmental profile. We decided to focus on organoids grown on the orbital plates to provide further characterization of the organoid generation pipeline, because the organoids grown in SpinΩ have been described elsewhere (Qian et al., 2016). We observed that 30-day organoids from orbital shaker cultures were positive for FOXG-1 (forebrain), PAX-6 (dorsal telencephalon), OTX-2 (retinal cells and midbrain), and islet-1 (hindbrain; Fig. 4a) showing diversification and development consistent with previous reports (Quadrato et al., 2017). We observed that, at 45 days, the organoids had pigmented regions (Fig. 4b, c), which were previously described to reproduce the formation of retinal pigmented epithelium (Quadrato et al., 2017). The pigmented regions were positive for the retinal cell marker glycogen synthetase (GS) (Fig. 4c). Electrophysiological recordings of 45-day organoids detected neural oscillations below 10 Hz in a pigmented organoid, but no firing was detected at the same developmental age in non-pigmented organoids (Supp. Fig. 5).

**Figure 4.**
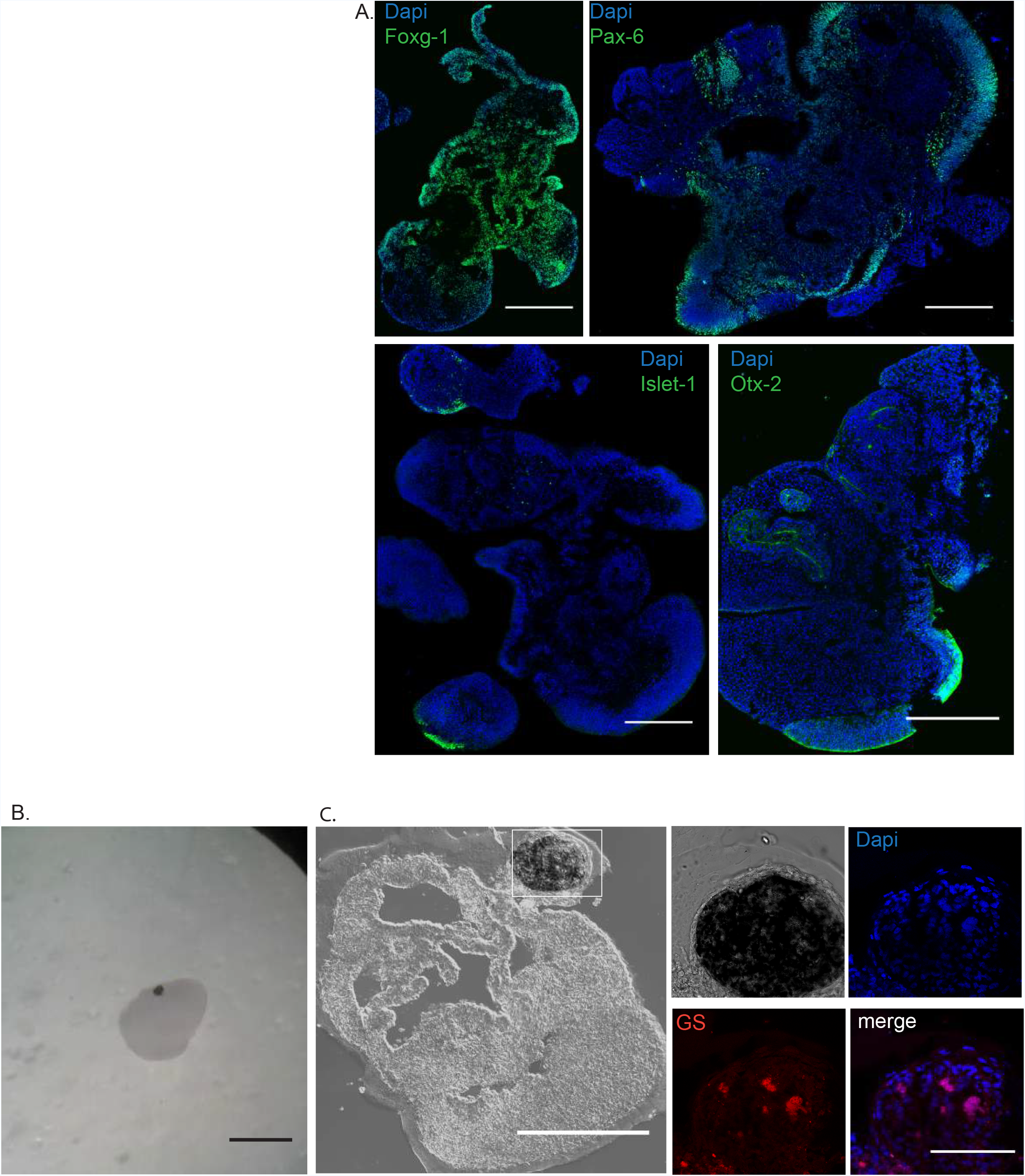
Cell types and brain regions represented in early brain organoid development. **a**. Immunostaining of 30-day-old organoids grown in the orbital shaker for FOXG-1, PAX-6, islet-1, and OTX-2. Bars = 500 μm; **b**. Stereoscopic image of an organoid with pigmented regions. Bar = 1 mm. **c**. Pigmented regions (box) of organoids after 45 days in culture. Bar = 1 mm. The pigmented regions (box) were positive for glycogen synthetase. Bar = 500 μm.

Proteomic analysis of organoids grown for 30 days led to the identification of 4,099 proteins (Supp. dataset). The 100 most abundant proteins (ordered by peptide spectral match) were analyzed by gene enrichment; 86 matched DAVID IDs (Huang et al., 2009a, 2009b). Gene enrichment analysis revealed that the majority (76.6%) of the identified proteins matched brain entries with high *p*-value. Proteins identified in our analysis are markers of the forebrain (BCL11B, DBI, CLU, SPARC), midbrain (OTX-2), and hindbrain (HOXA1). Retinal cell proteins were also identified: a general marker (GS) and those for the Muller glia (DKK3), photoreceptors (RCVRN), and retinal ganglion cells (NEFL). The appearance of these proteins at 30 days preceded the formation of the pigmented regions which were observed later, at 45 days (Fig. 4c).

## 4. Discussion

Brain organoids present cytoarchitecture that recapitulates brain tissue organization, offering a complex in vitro model for the study of brain normal and pathological development (Lancaster et al., 2013; Mariani et al., 2015; Paşca et al., 2015). Although brain organoid cultivation presents challenges related to the lack of reproducibility and scalability, we achieved high reproducibility of early-stage organoid size and growth by adding steps to a standard protocol (Lancaster and Knoblich, 2014b). This included the use of higher ROCKi concentrations and plate centrifugation in the EB formation step, which have been previously demonstrated to improve EB formation, but have never been applied to organoid formation protocol (Hookway et al., 2016; Horiguchi et al., 2014).

Scalability was achieved by applying two multiplex platforms: steering plates on an orbital shaker and the SpinΩ bioreactor (Qian et al., 2016). The SpinΩ was 3D printed according to the blueprints provided by Qian et al. (Qian et al., 2016). We encountered the following issues with SpinΩ use: 1) manual handling, as medium changes involved disassembly of a combination of pieces; 2) the need for sterilization for consecutive use; and 3) the maintenance of sterile conditions, as the equipment has 12 gears that could not be cleaned properly during the course of the experiment. These issues make the SpinΩ dependent on user skills, rendering it more prone to error and susceptible to contamination over the long timeframe of brain organoid cultivation (up to 8 months) (Lancaster and Knoblich, 2014b; Quadrato et al., 2017), when compared with the use of steering plates on an orbital shaker.

We suggest that the lower velocities of the SpinΩ, which affect nutrient mixing, may explain the decreased organoid growth seen on days 17 and 19, and the wide size distribution of organoids observed on day 30. Overall, our CFD analysis indicated that the fluid dynamic variables examined in this study are closer to those of the spinner bioreactor for the steering plate on the orbital shaker. Therefore, this method should be preferentially selected as a multiplex alternative to the use of a spinner bioreactor. The appearance of diverse brain regions and pigmented regions labelled with the retinal epithelium marker GS has been previously described (Quadrato et al., 2017) and related to a regional differentiation in organoids. Organoids produced using our protocol presented pigmented regions positive for GS suggesting that the technique described here may be appropriate for studies involving the complexity of early brain development.

In addition, proteomic analysis confirmed the organoids (produced in this study) show a protein profile that is compatible with several differentiated brain regions. Altogether, those results corroborate that the new proposed protocol opens a new widow, allowing the exploration, with multiple analyses, of important biomarkers of the morphological, genetic and molecular complexity of the human brain development under normal and abnormal conditions.

## Author contributions

L.G-S., N.M.E.A., F.T.M and S.K.R conceived and designed the study. L.G-S. and N.M.E.A. performed the experiments and data analysis. I.L.H., N.P.S, B.L., H.R.B.O. performed computational fluid dynamics analysis and interpretation. A.C.S. and S.R. performed electrophysiological measurements and analysis of the results. L.G-S., G.B.D and M.J. performed proteomics data collection and analysis. L.G-S. and N.M.E.A. prepared the figures and drafted the manuscript. H.R.B.O, S.R., F.T.M., and S.K.R. made critical revisions to the manuscript. All authors revised and approved the final manuscript.

## Acknowledgements

This work was supported by the Conselho Nacional de Pesquisa (CNPq), Coordenação de Aperfeiçoamento de Pessoal de Nível Superior (CAPES), Fundação Carlos Chagas Filho de Amparo à Pesquisa do Estado do Rio de Janeiro (FAPERJ), Banco Nacional de Desenvolvimento Econômico e Social (BNDES), Financiadora de Estudos e Projetos (Finep) and intramural grants from D’Or Institute for Research and Education (IDOR). We thank Marcelo Costa and Gabriela Vitória for their work in iPSC banking and expansion and Rodrigo Martins for kindly suppling antibodies for retinal cell identification. We also thank Bruna Palsen and Pítia Ledur for critical reading of the manuscript.

## Abbreviations

Three dimensional (3D)
human pluripotent stem cells (hPSC)
induced pluripotent stem cells (iPSC)
embryoid bodies (EBs)

## References

Bershteyn, M., Nowakowski, T.J., Pollen, A.A., Di Lullo, E., Nene, A., Wynshaw-Boris, A., and Kriegstein, A.R. (2017). Human iPSC-Derived Cerebral Organoids Model Cellular Features of Lissencephaly and Reveal Prolonged Mitosis of Outer Radial Glia. Cell Stem Cell 20, 435–449.e4.

Dakic, V., Minardi Nascimento, J., Costa Sartore, R., Maciel, R. de M., de Araujo, D.B., Ribeiro, S., Martins-de-Souza, D., and Rehen, S.K. (2017). Short term changes in the proteome of human cerebral organoids induced by 5-MeO-DMT. Sci. Rep. 7, 12863.

Eiraku, M., Takata, N., Ishibashi, H., Kawada, M., Sakakura, E., Okuda, S., Sekiguchi, K., Adachi, T., and Sasai, Y. (2011). Self-organizing optic-cup morphogenesis in three-dimensional culture. Nature 472, 51–56.

Fridley, K.M., Kinney, M.A., and McDevitt, T.C. (2012). Hydrodynamic modulation of pluripotent stem cells. Stem Cell Res. Ther. 3, 45.

Garcez, P.P., Loiola, E.C., Madeiro da Costa, R., Higa, L.M., Trindade, P., Delvecchio, R., Nascimento, J.M., Brindeiro, R., Tanuri, A., and Rehen, S.K. (2016). Zika virus impairs growth in human neurospheres and brain organoids. Science 352, 816–818.

Hookway, T.A., Butts, J.C., Lee, E., Tang, H., and McDevitt, T.C. (2016). Aggregate formation and suspension culture of human pluripotent stem cells and differentiated progeny. Methods 101, 11–20.

Horiguchi, A., Yazaki, K., Aoyagi, M., Ohnuki, Y., and Kurosawa, H. (2014). Effective Rho-associated protein kinase inhibitor treatment to dissociate human iPS cells for suspension culture to form embryoid body-like cell aggregates. J. Biosci. Bioeng. 118, 588–592.

Huang, D.W., Sherman, B.T., and Lempicki, R.A. (2009a). Bioinformatics enrichment tools: paths toward the comprehensive functional analysis of large gene lists. Nucleic Acids Res. 37, 1–13.

Huang, D.W., Sherman, B.T., and Lempicki, R.A. (2009b). Systematic and integrative analysis of large gene lists using DAVID bioinformatics resources. Nat. Protoc. 4, 44–57.

Lancaster, M. a., and Knoblich, J. a. (2014a). Organogenesis in a dish: modeling development and disease using organoid technologies. Science 345, 1247125.

Lancaster, M.A., and Knoblich, J.A. (2014b). Generation of cerebral organoids from human pluripotent stem cells. Nat. Protoc. 9, 2329–2340.

Lancaster, M.A., Renner, M., Martin, C.-A., Wenzel, D., Bicknell, L.S., Hurles, M.E., Homfray, T., Penninger, J.M., Jackson, A.P., and Knoblich, J.A. (2013). Cerebral organoids model human brain development and microcephaly. Nature 501, 373–379.

Lancaster, M.A., Corsini, N.S., Wolfinger, S., Gustafson, E.H., Phillips, A.W., Burkard, T.R., Otani, T., Livesey, F.J., and Knoblich, J.A. (2017). Guided self-organization and cortical plate formation in human brain organoids. Nat. Biotechnol. 35, 659–666.

Mariani, J., Coppola, G., Zhang, P., Abyzov, A., Provini, L., Tomasini, L., Amenduni, M., Szekely, A., Palejev, D., Wilson, M., et al. (2015). FOXG1-Dependent Dysregulation of GABA/Glutamate Neuron Differentiation in Autism Spectrum Disorders. Cell 162, 375–390.

Monzel, A.S., Smits, L.M., Hemmer, K., Hachi, S., Moreno, E.L., van Wuellen, T., Jarazo, J., Walter, J., Brüggemann, I., Boussaad, I., et al. (2017). Derivation of Human Midbrain-Specific Organoids from Neuroepithelial Stem Cells. Stem Cell Rep. 8, 1144–1154.

Murillo, J.R., Goto-Silva, L., Sánchez, A., Nogueira, F.C.S., Domont, G.B., and Junqueira, M. (2017). Quantitative proteomic analysis identifies proteins and pathways related to neuronal development in differentiated SH-SY5Y neuroblastoma cells. EuPA Open Proteomics 16, 1–11.

Nampe, D., Joshi, R., Keller, K., and Nieden, N.I. (2017). Impact of Fluidic Agitation on Human Pluripotent Stem Cells in Stirred Suspension Culture.

Paşca, A.M., Sloan, S.A., Clarke, L.E., Tian, Y., Makinson, C.D., Huber, N., Kim, C.H., Park, J.-Y., O’Rourke, N.A., Nguyen, K.D., et al. (2015). Functional cortical neurons and astrocytes from human pluripotent stem cells in 3D culture. Nat. Methods 12, 671–678.

Qian, X., Nguyen, H.N., Song, M.M., Hadiono, C., Ogden, S.C., Hammack, C., Yao, B., Hamersky, G.R., Jacob, F., Zhong, C., et al. (2016). Brain-Region-Specific Organoids Using Mini-bioreactors for Modeling ZIKV Exposure. Cell 165, 1238–1254.

Quadrato, G., Nguyen, T., Macosko, E.Z., Sherwood, J.L., Min Yang, S., Berger, D.R., Maria, N., Scholvin, J., Goldman, M., Kinney, J.P., et al. (2017). Cell diversity and network dynamics in photosensitive human brain organoids. Nature 545, 48–53.

Sartore, R.C., Cardoso, S.C., Lages, Y.V.M., Paraguassu, J.M., Stelling, M.P., Madeiro da Costa, R.F., Guimaraes, M.Z., Pérez, C.A., and Rehen, S.K. (2017). Trace elements during primordial plexiform network formation in human cerebral organoids. PeerJ 5, e2927.

Wang, Y., Chou, B.-K., Dowey, S., He, C., Gerecht, S., and Cheng, L. (2013). Scalable expansion of human induced pluripotent stem cells in the defined xeno-free E8 medium under adherent and suspension culture conditions. Stem Cell Res. 11, 1103–1116.

